# Automated Removal of Corrupted Tilts in Cryo-Electron Tomography

**DOI:** 10.1101/2025.03.13.642992

**Authors:** Tomáš Majtner, Beata Turoňová

## Abstract

Cryo-electron tomography (cryo-ET) enables the visualization of macromolecular structures in their near-native cellular environment. However, acquired tilt series are often compromised by image corruption due to drift, contamination, and ice reflections. Manually identifying and removing corrupted tilts is subjective and time-consuming, making an automated approach necessary. In this study, we present a deep learning-based method for automatically removing corrupted tilts. We evaluated 13 different neural network architectures, including convolutional neural networks (CNNs) and transformers. Using a dataset of 435 annotated tilt series, we trained models for both binary and multiclass classification of corrupted tilts. We demonstrate the high efficiency and reliability of these automated approaches for removing corrupted tilts in cryo-ET and provide a framework, including models trained on cryo-ET data, that allows users to apply these models directly to their tilt series, improving the quality and consistency of downstream cryo-ET data processing.

## 1 Introduction

Cryo-electron tomography (cryo-ET) is a powerful imaging technique that enables the visualization of macromolecular structures in their native cellular context. Data acquisition in cryo-ET involves the collection of tilt series (TS), where individual images (tilts) capture a specimen at varying angles. Several factors can compromise image quality in TS. For example, large electron-dense objects, such as grid bars, lamella edges, or ice contamination, may partially or completely obstruct the electron beam, resulting in large black areas in tilt images. Specimen drift can introduce image blurring, while sub-vitreous specimens may produce ice reflections that interfere with tilt alignment. These artifacts complicate subsequent processing steps, including TS alignment and contrast transfer function (CTF) estimation, ultimately resulting in suboptimal tomogram reconstruction. The TS alignment is especially crucial for high-resolution structure determination using subtomogram averaging as demonstrated by Turoňová et al.^1^. Supplementary Figure 1 provides a visual illustration of the effect of cleaning on tomogram reconstruction, along with a summary of the residual mean values for tilt series alignment before and after cleaning. For this reason, it is common practice to identify and exclude problematic tilts early in the cryo-ET workflow^2–8^.

Currently, identifying and removing sub-optimal tilts is typically performed manually, relying on the subjective assessment of individual users. This process is not only time-consuming and tedious but also highly variable, as different users may apply inconsistent criteria when evaluating tilt quality. For large PACE datasets^9^, even with user-friendly GUI tools, manual tilt cleaning can take several minutes per TS, making it a major bottleneck in the workflow. Despite these challenges, efforts toward automated tilt removal remain limited. The *Tomo Live* software^10^ includes the Stack Cleaner module, which calculates the mean and standard deviation of correlation values from the correlation matrix. For each tilt image, a check is performed to assess whether its histogram is consistent with those of other tilt images. Similarly, AreTomo^11^ includes a feature that can exclude certain tilts, particularly those that are predominantly black due to obstructions. However, this approach remains relatively basic and requires a fixed threshold parameter and consequently additional manual post-processing by the user.

Manual inspection remains standard practice, as seen in RELION-5 pipeline^5^, where *Exclude tilt-images* job launches a Napari-based^12^ viewer that displays the images in a tilt series and allows the user to select unsuitable images. Other examples of manual exclusion are in recent studies on nuclear pore permeability^3^ or on the palisade layer of the poxvirus core^4^, where corrupted projections were removed after visual inspection. The manual inspection of TS is a bottleneck on cryo-ET workflows, especially as cryo-ET datasets grow larger due to advances in automation and throughput^9^ of modern electron microscopes^13^. This underscores the need for an automated approach to tilt removal that is efficient and objective across the users.

In recent years, machine learning techniques have been introduced to enhance data analysis in cryo-ET^14, 15^. For instance, tools like cryoCARE^16^ demonstrated the feasibility of deep learning for tasks such as denoising. These approaches leverage convolutional neural networks (CNNs), which are trained on large annotated datasets, to automate previously manual tasks with high accuracy and reproducibility. Although these methods do not address tilt stack filtering directly, they provide a framework for applying similar techniques to the tilt removal problem.

While the methods described above represent notable advancements, a fully automated and objective solution for corrupted tilt removal has yet to be introduced. In this study, we conducted a comprehensive evaluation of 13 deep learning models with varying learning paradigms and architectures. Our results demonstrate that Swin transformers achieve an area under the ROC curve (AUC) exceeding 0.96 and an F1-score above 0.94, indicating a strong correlation with manual annotations. This enables a consistent evaluation of tilt quality, addressing both subjectivity and speed limitations. By leveraging advanced neural networks trained on annotated tilt series, we can now quickly identify and exclude problematic tilts, allowing researchers to focus on downstream analysis.

## 2 Workflow and Datasets

We tested 13 deep learning models using 435 tilt series from six different datasets. The tilt series were provided by their respective authors, all experienced cryo-ET users, including their manual annotations. The models were trained to simulate the process of categorizing tilt images as corrupted or good. The overview of our method is depicted in Figure 1.

**Figure 1.**
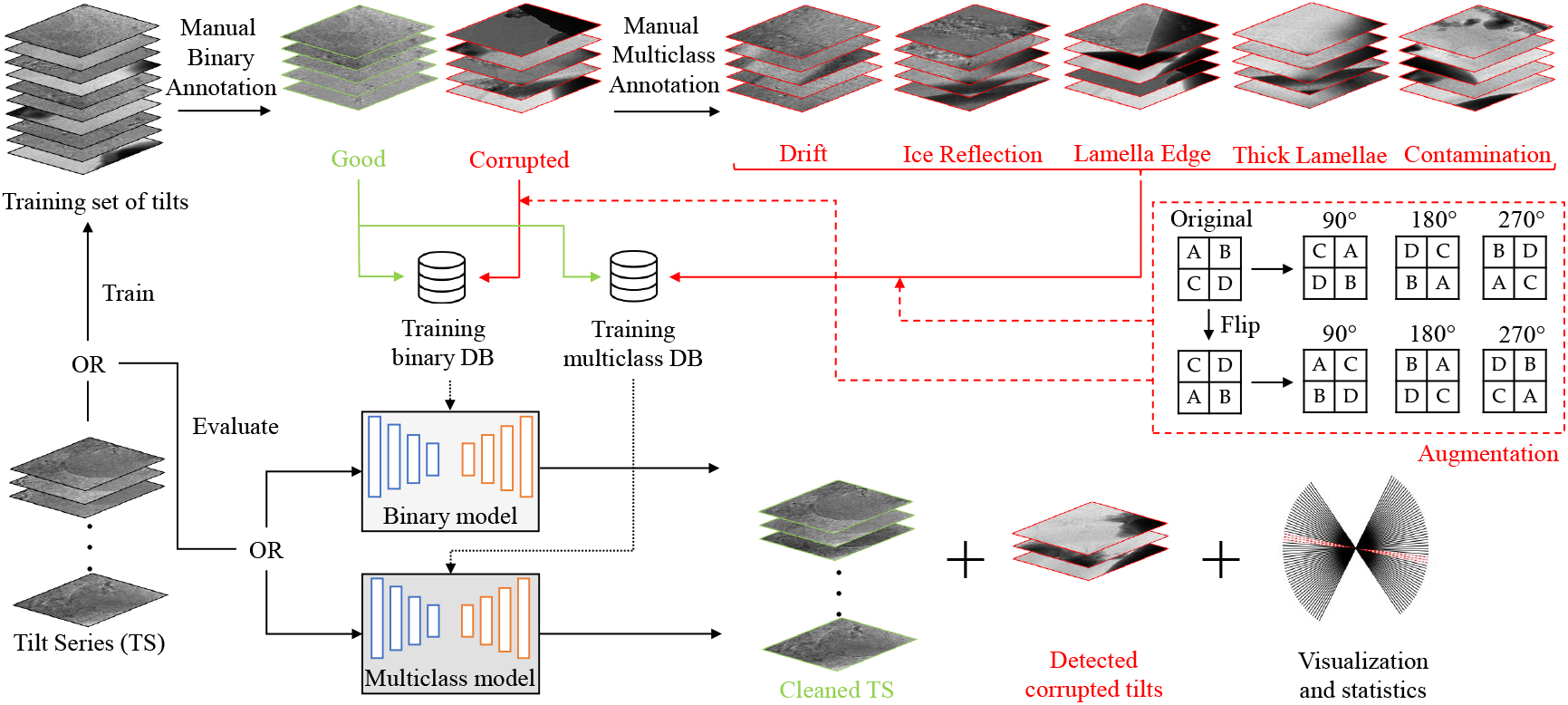
Workflow. The training set of tilt images was manually categorized into corrupted and good tilts. For multiclass classification, the corrupted tilts were further divided into five categories based on the specific type of image artifact. Since the number of corrupted tilts was substantially lower than that of good tilts, data augmentation was applied to balance the dataset. The training datasets were then used to develop models for both binary and multiclass classification. During evaluation, a user can input a tilt series and select either a binary or multiclass model. The selected model will then provide a cleaned tilt series, a list of excluded tilts, and a visualization of the removed tilts.

Six *in situ* datasets from lamellae were used in this study, each annotated by a different user who acquired the data. These datasets were selected because lamellae samples are typically more challenging to process and, consequently, more difficult to clean. We refer to these datasets as D1 through D6 and provide an overview in Table 1.

**Table 1.** Overview of employed datasets. Each dataset is from a different user and we summarize here the number of TS, cell type, pixel size, and tilt increment used in each collection. In D4, 26 TS have 1°, 41 TS have 2°, and 58 TS have 3° tilt increment.

| Dataset | # of TS | Cell Type | Pixel Size (Å/px) | Tilt (°) |
| --- | --- | --- | --- | --- |
| D1 | 23 | Human Embryonic Kidney (HEK Flp-In T-Rex) | 1.971 | 2 |
| D2 | 108 | Human macrophages | 2.414 | 2 |
| D3 | 11 | Human skin fibroblasts | 2.414 | 2 |
| D4 | 125 | *Dictyostelium discoideum* | 1.971 | 1, 2, 3 |
| D5 | 145 | *Dictyostelium discoideum* | 2.18 | 2 |
| D6 | 23 | Human Embryonic Kidney (HEK Flp-In T-Rex) | 2.682 | 2 |

The analyzed TS were divided into training (315 TS), validation (29 TS), and test (91 TS) subsets. Each tilt series contained at least one tilt image identified by the original user as a corrupted tilt. The detailed distributions of corrupted and good tilts, along with their respective division into training, validation, and test subsets, are summarized in Table 2.

**Table 2.**
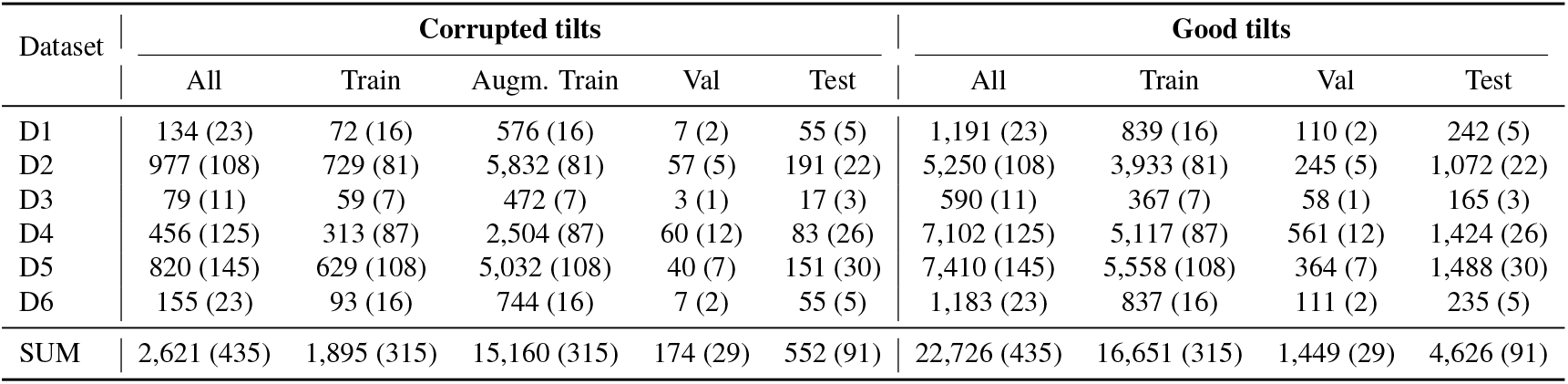
Dataset statistics for corrupted and good tilts. Corrupted tilts were augmented using horizontal flipping and rotations of 90°, 180°, and 270° to balance the training portion of the dataset. To avoid potential bias during training, all tilts from a single tilt series are included exclusively in either the training, validation, or testing portion of the dataset. The values in the table specify the total number of tilt images in each category, with the corresponding number of tilt series indicated in parentheses.

Each tilt series contained a larger number of good tilts compared to corrupted ones and thus there was a marked imbalance between the total number of images in the two categories (2,621 corrupted tilts vs. 22,726 good tilts). While this ratio reflects the realistic distribution in the validation and test subsets, the imbalance posed a challenge for model training. To address this, we employed an image augmentation strategy that included horizontal flipping and rotations of exactly 90°, 180°, and 270° to avoid introducing interpolation artifacts in data. This process, illustrated also in Figure 1, effectively increased the size of the corrupted training dataset by a factor of eight.

To further advance this approach, we manually classified a subset of 1,242 corrupted tilt images from datasets D1–D4, categorizing them into five distinct classes: contamination (CO), drift (DR), ice reflection (IR), lamella edge (LE), and thick lamellae (TL). While most of these categories have clear definitions, the thick lamellae category specifically refers to images taken at high tilt angles, where the effective specimen thickness increases due to projection geometry. Although the physical thickness of the lamella remains constant across all tilts in a series, higher tilt angles cause greater material overlap since the path of the electron beam within the specimen gets longer, leading to reduced image contrast and visibility. In some instances, this effect is further exacerbated by milling artifacts, such as curtaining, which may introduce dark regions that persist across all tilts. Thus, the classification of thick lamellae focuses on identifying images where these factors markedly impair visibility, making them more challenging for downstream processing. The main advantage of multiclass labeling is that users can selectively remove any specified categories of corrupted tilts.

These corrupted tilts, combined with 14,473 good ones from the same tilt series, formed a second large, annotated dataset with multiclass labels that was subsequently split into training, validation, and test subsets (see Table 3 for details). We then trained the same collection of deep learning architectures on this multiclass dataset, enabling the models not only to identify whether a tilt image is corrupted, but also to specify the reason for exclusion. Tilts originally excluded due to defocus values were omitted from the multiclass dataset, as they can be easily identified and removed during post-processing using defocus estimation algorithms such as Gctf^17^, CTFFind4^18^ or CTFFind5^19^, which users run as part of the tomogram reconstruction workflow.

**Table 3.** Data distribution across different classes for train, augmented train, validation, and test sets. OK is good tilt, DR is drift, IR is ice reflection, LE is lamella edge, TL is thick lamellae, and CO is contamination.

|  | OK | DR | IR | LE | TL | CO |
| --- | --- | --- | --- | --- | --- | --- |
| Train | 10,506 | 44 | 124 | 146 | 224 | 320 |
| Augmented Train | 10,506 | 352 | 992 | 1,168 | 1,792 | 2,560 |
| Validation | 757 | 7 | 19 | 4 | 25 | 13 |
| Test | 3,210 | 28 | 51 | 65 | 46 | 126 |

## 3 Methods Overview

In this study, we examined conventional convolutional neural networks (CNNs) and transformers. Both approaches are complementary and fall under the umbrella of deep learning. The key differences are that CNNs use pixel-based data handling to capture local spatial features via convolutions. They have lower computational cost and are more interpretable due to localized filters. On the other hand, transformers use patch-based data handling to capture global relationships via self-attention. They have high computational and memory cost but scale effectively with larger datasets^20–22^.

In addition to CNNs and transformers, we also explored non-deep-learning approaches, such as entropy-based feature extraction combined with logistic regression. However, these traditional methods did not achieve performance levels comparable to the deep learning models and we have not included them in this study.

### 3.1 CNNs

From the CNNs category, we selected EfficientNet^23^, MobileNet_v2^24^, and ResNet^25^. EfficientNet family of CNNs uses a compound scaling method that uniformly scales depth, width, and resolution to achieve optimal performance across various model sizes. In particular, we chose lightweight EfficientNet_b0, together with scaled up EfficientNet_b3, and EfficientNet_b7, which is one of the largest models in the EfficientNet family, with considerably increased capacity.

MobileNet_v2 is a lightweight convolutional neural network that introduces inverted residual blocks and linear bottlenecks to reduce computational cost. The model is widely used in applications requiring real-time inference. ResNet (Residual Network) is a commonly used deep learning architecture that introduced residual connections, enabling very deep networks to avoid vanishing gradients and improve training stability. From the ResNet family, we employed ResNet-18, ResNet-50, and ResNet-101, where the number reflects the depth of the network. More technical details about each network can be found in their respective cited articles mentioned above.

### 3.2 Transformers

Transformer models leverage a mechanism called self-attention^22^, allowing them to process and weigh relationships between all parts of an image simultaneously, rather than focusing on small, localized regions like CNNs. Images are divided into patches, which are flattened and linearly embedded into a sequence of vectors, where each patch represents a part of the image. Transformers excel when pre-trained on massive datasets and fine-tuned for specific tasks.

We considered four transformer-based architectures. Vision transformer (ViT) introduced by Dosovitskiy et al.^26^ applies transformer architectures directly to image recognition tasks by dividing images into patches and processing them as sequences. ViT was the first to successfully apply transformer architecture to vision tasks. Its key feature includes patch embedding, where the input image is divided into fixed-size patches and each patch is flattened into a 1D vector. These vectors are linearly embedded into a higher-dimensional space to serve as input tokens.

Data-efficient image transformer (DeiT), developed by Touvron et al.^27^, enhances ViT by incorporating training strategies that reduce data requirements, enabling effective training without extensive datasets. DeiT uses techniques such as strong data augmentation, regularization, and knowledge distillation to train on ImageNet without pre-training on larger datasets.

Swin transformer, proposed by Liu et al.^28^, was designed to address the limitations of ViT in processing high-resolution images efficiently. Instead of applying global attention, Swin restricts attention to local non-overlapping windows. From the Swin family, we tested Swin_tiny, a small version of the Swin transformer, designed for lightweight applications with lower computational cost; Swin_base, a balanced version with more parameters than Swin_tiny, offering a trade-off between efficiency and accuracy, and Swin_large, the largest version, featuring the highest number of parameters and computational complexity, achieving state-of-the-art performance in vision tasks.

Lastly, BEiTv2 (Bidirectional Encoder representation from Image Transformers v2) was presented by Peng et al.^29^. It advances self-supervised learning for vision transformers by employing a semantic-rich visual tokenizer that encodes images into tokens. These tokens provide a better initialization for the transformer, improving its downstream performance. For more technical details on each network we refer the reader to the respective cited article for each method.

### 3.3 Training Parameters

All networks use initial weights from the ImageNet dataset^30^, and fine-tuning, that is commonly referred to as transfer learning^31^, was applied to adapt them for corrupted tilt recognition. Specifically, for all tested networks, the original classification layer was replaced with a new layer tailored to our task: a binary classification layer for models with two classes or a multiclass layer for models with six classes.

For fine-tuning, we employed the Adam optimizer with the learning rate of 0.0001 and a mini-batch size of 32 images. As for the loss function, we used cross-entropy loss. Each network was trained for 50 epochs (see Supplementary Figure 2 for the effect of epochs number on the F1 scores), with the best-performing model selected based on the lowest validation loss across all iterations. The input images were resized to 224 *×* 224 pixels, except for EfficientNet_b3 and EfficientNet_b7, where an input size of 320 *×* 320 pixels was used. Training and validation were conducted using PyTorch with our custom-developed code, available in the Code Availability section.

During training, performance was monitored using a validation dataset, and the final version of each model was evaluated on an independent test dataset. All tests were performed on a machine with AMD EPYC 7543P 32-Core Processor and two NVIDIA A100 80 GB GPU cards. Training time, as well as evaluation, can vary greatly among different models. (see Table 4).

**Table 4.**
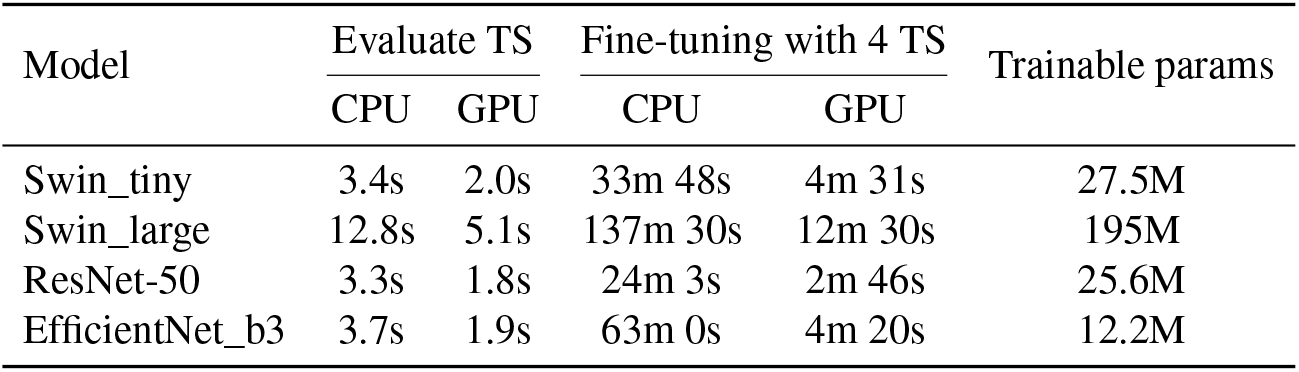
Summary of running times and network sizes. For tilt series evaluation, we used a single TS with 50 tilt images. To test the fine-tuning times, we used four random TS from the D4 dataset.

### 3.4 Evaluation Metrics

To evaluate the performance of our models, we employed the F1-score, the Receiver Operating Characteristic (ROC) curve, and the Area Under the Curve (AUC-ROC). The F1-score is a widely used metric in classification tasks, particularly for imbalanced datasets, as it balances precision and recall. It is defined as the harmonic mean of precision and recall:

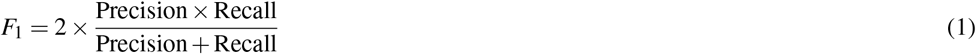

where precision and recall are given by:

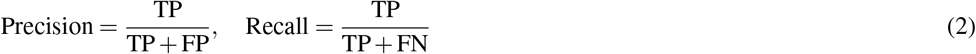

Here, TP, FP, and FN represent true positives, false positives, and false negatives, respectively.

The ROC curve is a graphical representation used to assess the performance of a binary classifier by plotting the true positive rate (TPR) against the false positive rate (FPR) at various threshold settings. These rates are defined as:

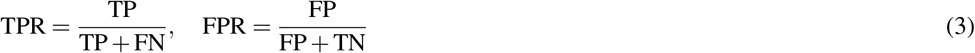

where TN denotes true negatives.

The AUC-ROC quantifies the classifier’s ability to discriminate between positive and negative classes. A higher AUC value (closer to 1) indicates a better-performing model, while a value of 0.5 suggests random classification.

For multiclass classification problems, the ROC curve needs to be adapted and we used One-vs-Rest (OvR) strategy. In the OvR approach, an ROC curve is generated for each class by treating that class as the positive class and all other classes as the negative class. For each class *c*, the TPR and FPR are computed as:

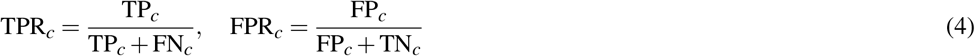

where *T P*_*c*_, *FP*_*c*_, *TN*_*c*_, and *FN*_*c*_ represent the number of true positives, false positives, true negatives, and false negatives for class *c*, respectively.

The AUC is then computed for each class separately:

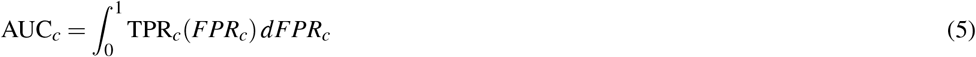

The final multiclass AUC score is in our case averaged across all classes using non-weighted mean commonly known as macro-averaging.

## 4 Results

### 4.1 Performance of Binary Classification Models

Figure 2 presents the F1-scores for the evaluated binary classification models across different training strategies, where the individual black dots are the individual datasets. Models trained using all six augmented training parts of D1-D6 (*Train_all*) exhibit superior performance compared to those employing leave-one-out cross-validation (*Train_5DS*). In the latter strategy, we always removed one training dataset and evaluated on the corresponding test dataset (e.g. train on D1-D5, evaluate on D6). This suggests that there are certain variations between datasets and that each user has a slightly different subjective criteria for removing corrupted tilts. The difference is not very large but highlights the need for a unified approach. We also tested dataset specific training (*Train_1DS*), where we always trained and evaluated each dataset separately. We can observe much higher dispersion of F1-score values for certain networks in these tests, which suggests that not all methods are suitable for the training with lower numbers of samples. Overall, transformer-based models, particularly Swin transformers, consistently achieve higher F1-scores than CNN-based models.

**Figure 2.**
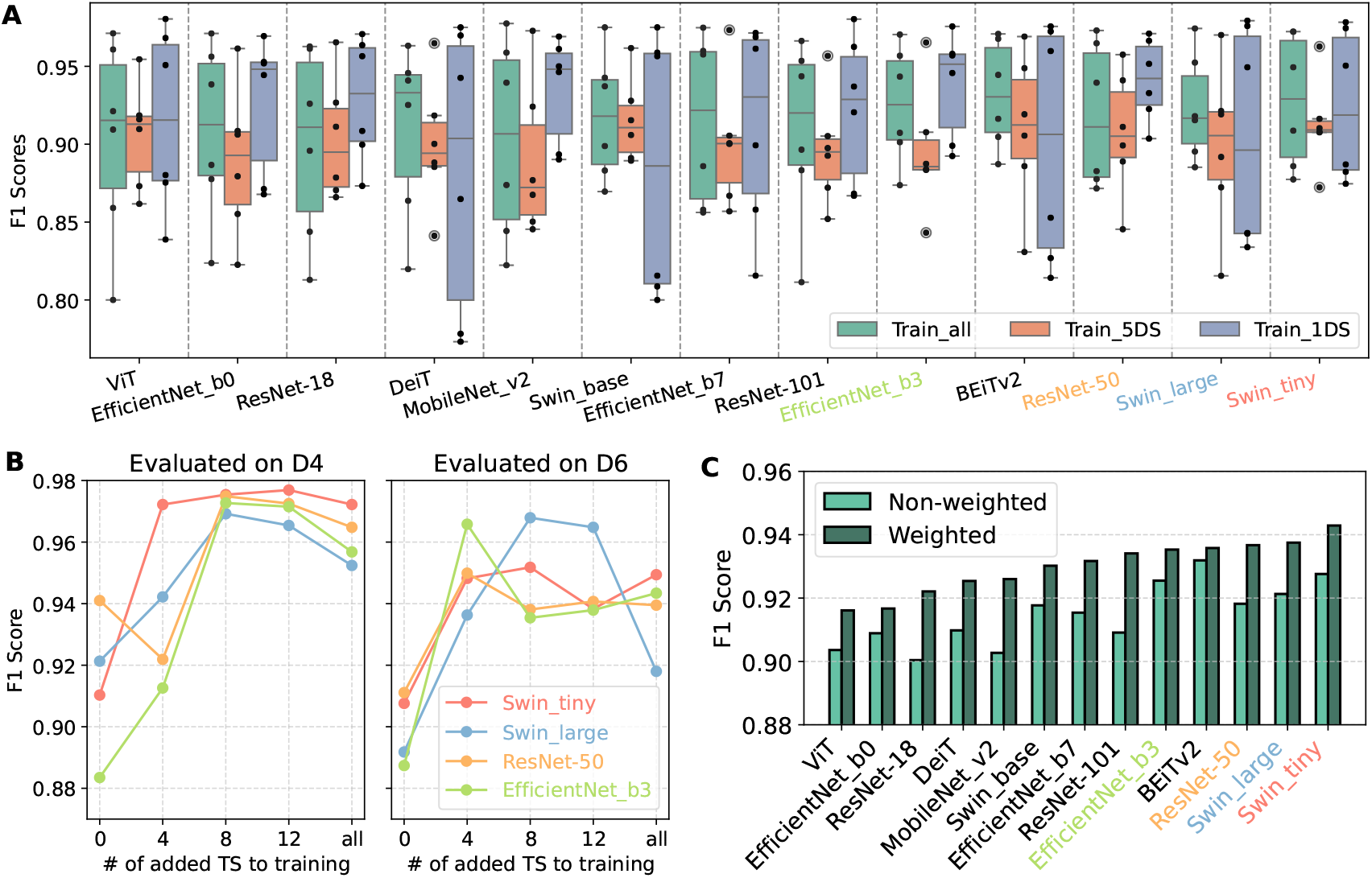
(A) Binary models ordered by weighted F1-score. Train_all uses all training samples, train_5DS employs a leave-one-out cross-validation strategy, and train_1DS consists of separately trained models using the training portion of D1–D6, with evaluation performed on the corresponding test portion. (B) Fine-tuning of pre-trained networks with the limited number of samples. After adding only four TS from training part of the dataset, the F1 score increases for all models. (C) The summary of weighted and non-weighted F1-scores for all tested models.

A key advantage of pre-trained deep learning models is their ability to generalize with limited training data. To assess if this also applies to our TS cleaning, we selected four fine-tuned models (two best performing CNNs and two best performing transformers, all highlighted with a specific color in Figure 2) with a progressively increasing number of training tilt series. Figure 2B shows that after adding just four additional training tilt series, all models exhibit a notable improvement in F1-score. This trend suggests that transfer learning enables models to rapidly adapt to tilt classification tasks, making it particularly beneficial for users with limited annotated datasets. By leveraging a pre-trained network, users can efficiently tailor the model to their specific cleaning strategy without requiring extensive manual labeling. We recommend applying this approach not only to *in situ* samples but also to in-vitro specimens or non-milled cells, where users can easily adapt our provided networks to their specific data, even if it differs from in-situ conditions. Figure 2C provides the summary of weighted and non-weighted F1-scores for all tested models. In the weighted scenario, we take into account the number of tilt images in each tested dataset. The higher difference between weighted and non-weighted values suggests that the network performs better with the higher number of training samples.

### 4.2 ROC Curves and Missclassified Tilts

The classification performance of binary and multiclass models was further evaluated using ROC curves and AUC-ROC scores (Figure 3A). For binary classification, EfficientNet_b3 achieves the highest AUC (0.9049), followed closely by ResNet-50 (0.8945) and Swin-large (0.8636). Among transformer-based models, Swin-large and Swin-tiny perform comparably, but their AUC values remain slightly below those of the top CNNs.

**Figure 3.**
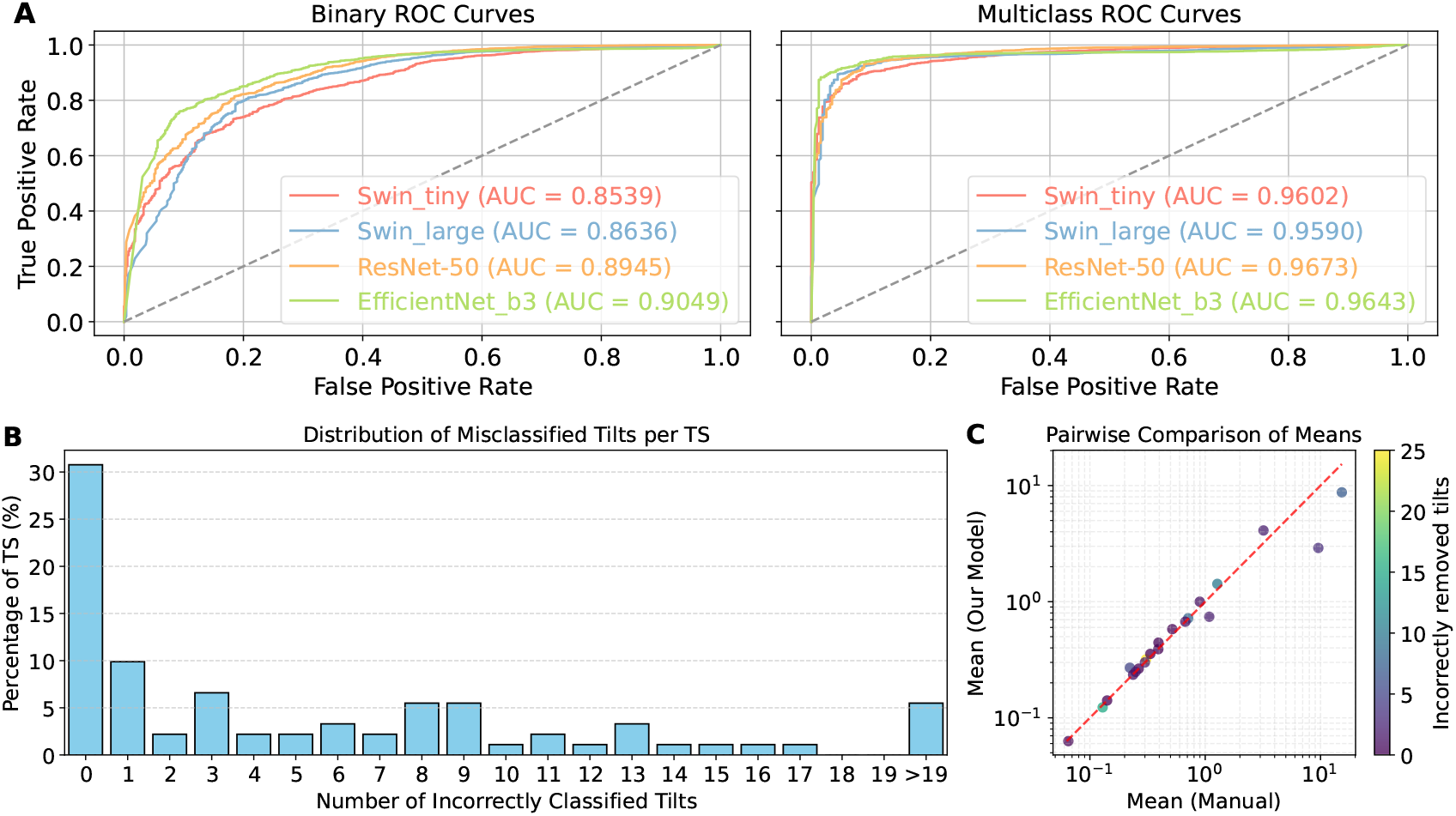
(A) ROC curves for the two best-performing CNNs and the two best-performing transformers, along with their corresponding AUC-ROC values. (B) Distribution of misclassified tilts per TS for Swin_large transformer. (C) Pairwise comparison of residual means from tilt series alignment between manual cleaning and automated cleaning with Swin_large transformer for the D4 dataset, showing a high correlation.

For multiclass classification, ResNet-50 achieves the highest AUC score of 0.9673, indicating strong performance in distinguishing between different types of tilt artifacts. EfficientNet_b3 (AUC = 0.9643) and Swin-large (AUC = 0.9590) also demonstrate excellent generalization across multiple artifact categories. The consistent superiority of ResNet-50 and EfficientNet_b3 suggests that CNNs remain highly competitive in both binary and multiclass settings, despite the advantages of transformer models in feature extraction. While the F1-score is most suitable for imbalanced datasets where false positives and false negatives are critical, a higher AUC indicates that the model more effectively ranks positive instances above negative ones. These results suggest that all models excel in recognizing good tilts, while CNNs are more effective at identifying corrupted tilts but may have slightly lower accuracy in detecting good ones.

To quantitatively assess the deviation between automatic and manual cleaning at the TS level, we analyzed the classification results across 91 test TS. The distribution of misclassified tilts per TS is presented in Figure 3B. Our findings indicate that in over 30% of TS, the same tilts were removed by both manual cleaning and our Swin_large transformer model. Furthermore, the majority of TS exhibit only minor discrepancies, with fewer than 5% of TS containing 20 or more tilts misclassified relative to manual cleaning. These differences primarily arise in cases where the deep learning model suggests discarding a substantial number of tilts, whereas human users may opt to retain them for further processing. Importantly, we provide an interactive visualization tool that displays all misclassified images (see Supplementary Figure 3), allowing users to review and, if necessary, override the automated cleaning decisions.

We also analyzed the mean residuals from the tilt series alignment as performed in eTomo^32^ using TS cleaned manually and automatically (Figure 3C). Our evaluation on test samples from the D4 dataset revealed that residual means remain largely similar, even in cases where multiple tilts were incorrectly removed. Specifically, 14 data points lie above the dashed red line, indicating higher residuals for automatic cleaning compared to manual cleaning, with a mean difference of only 0.0686 and a standard deviation of 0.1673. Conversely, 12 data points fall below the line, with a mean difference of 0.8006 and a standard deviation of 1.8126. These findings demonstrate a strong correlation in residual mean values, even when a higher number of tilts were misclassified relative to manual annotation. Visual examples of misclassified images are illustrated in Supplementary Figure 13.

### 4.3 Confusion Matrices and Error Analysis

To assess the performance of our top-performing models, we first analyzed confusion matrices for Swin-tiny, Swin-large, ResNet-50, and EfficientNet_b3 (Figure 4). While all multiclass models perform well in identifying good tilts, their performance varies across different artifact categories. Contamination and lamella edges are typically easy to recognize, resulting in high accuracy. In contrast, ice reflection artifacts and thick lamellae are slightly more challenging to detect. In practice, ice reflection artifacts become most apparent when scrolling through a tilt series, as they appear as dark spots with white halos that are absent in the previous and subsequent tilts. However, since tilts are treated independently in this analysis, these artifacts may instead resemble subtle dark blobs, which are common in tilt images and thus harder to distinguish. The most challenging artifact, however, is drift, which was poorly recognized, particularly by transformer models. This may be due to the low resolution at which the networks process data, as well as the limited number of drift samples available. These findings suggest that while deep learning models are highly effective in automated tilt removal, certain artifact types remain difficult to classify.

**Figure 4.**
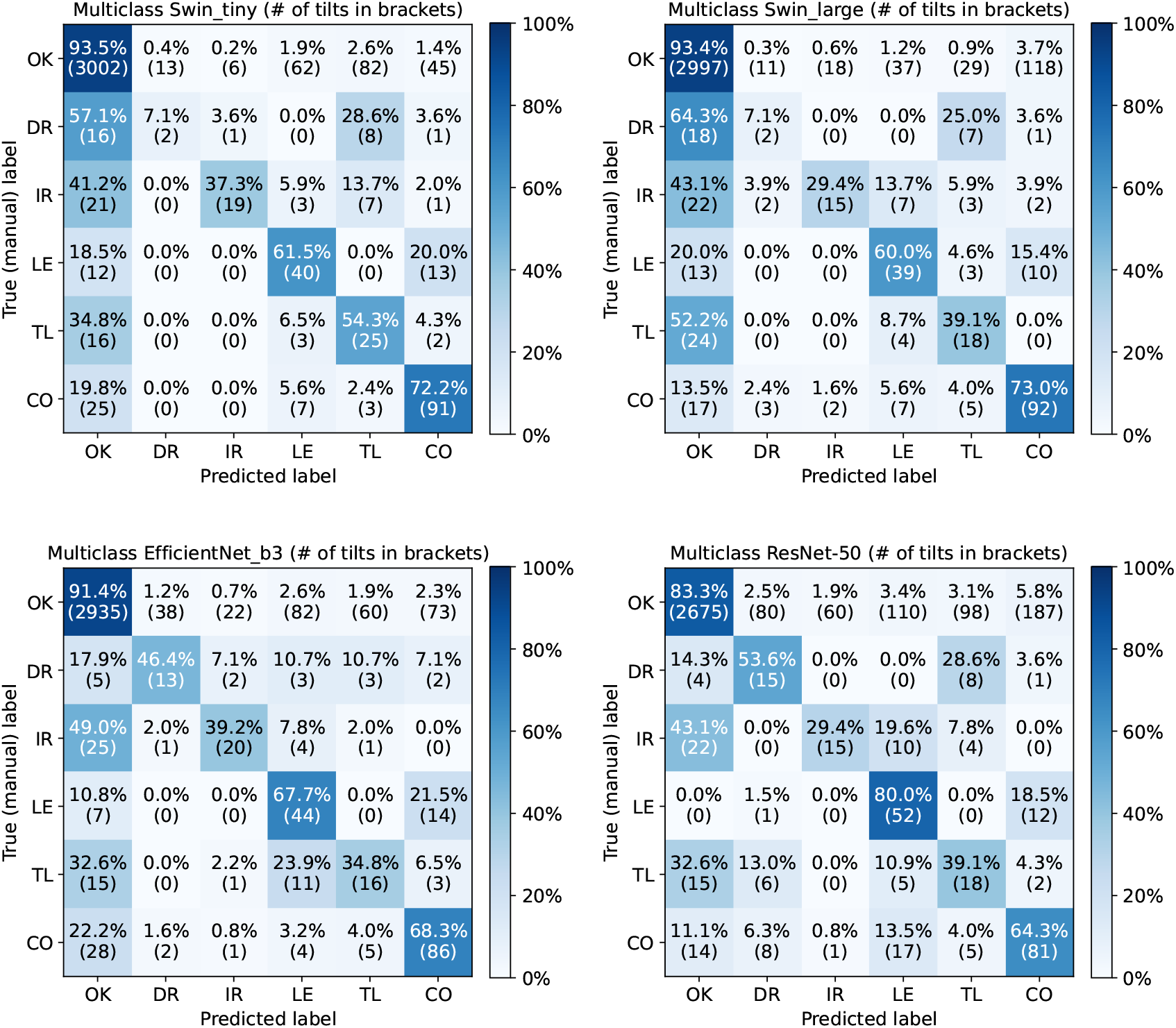
Confusion matrices for Swin-tiny, Swin-large, ResNet-50, and EfficientNet_b3, where OK is good tilt, DR is drift, IR is ice reflection, LE is lamella edge, TL is thick lamellae, and CO is contamination. All models excel at identifying corrupted tilts but the performance varies across different artifact categories. The results for remaining trained models are in Supplementary Figures 4,5,6,7,8,9,10,11,12.

### 4.4 Binary Evaluation of Multiclass Models

In addition to the multiclass analysis, we also evaluated the models using a binary classification approach, grouping predictions into corrupted and good tilts. This evaluation provides a simplified perspective on the models’ ability to distinguish between good and corrupted tilts, complementing the detailed multiclass results. Table 5 presents these binary evaluation results. Among the tested models, Swin-large achieves the best performance, correctly classifying 93.4% of good tilts and 70.3% of corrupted tilts. EfficientNet_b3 and ResNet-50 also exhibit high accuracy in detecting good tilts but show reduced sensitivity to corrupted tilts, with misclassification rates of 8.6% and 16.7%, respectively.

**Table 5.**
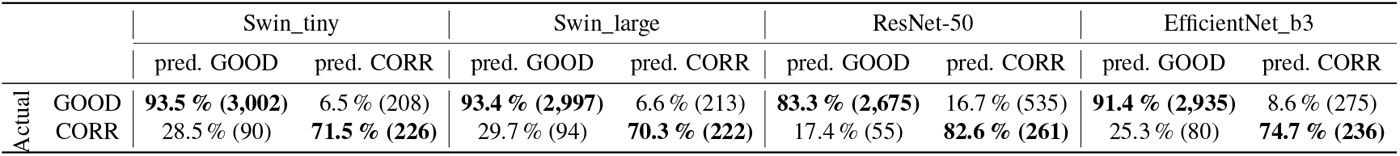
Binary evaluation of multiclass classification for two best performing CNNs and two best performing transformers. The results for remaining trained models are in Supplementary Figures 4,5,6,7,8,9,10,11,12.

|  |  | Swin_tiny |  | Swin_large |  | ResNet-50 |  | EfficientNet_b3 |  |
| --- | --- | --- | --- | --- | --- | --- | --- | --- | --- |
|  |  | pred. GOOD | pred. CORR | pred. GOOD | pred. CORR | pred. GOOD | pred. CORR | pred. GOOD | pred. CORR |
| Actual | GOOD | 93.5 % (3,002) | 6.5 % (208) | 93.4 % (2,997) | 6.6 % (213) | 83.3 % (2,675) | 16.7 % (535) | 91.4 % (2,935) | 8.6 % (275) |
|  | CORR | 28.5 % (90) | 71.5 % (226) | 29.7 % (94) | 70.3 % (222) | 17.4 % (55) | 82.6 % (261) | 25.3 % (80) | 74.7 % (236) |

Interestingly, CNN-based models, such as ResNet-50, demonstrate high precision but lower recall for corrupted tilts, implying that they are conservative in labeling tilts as being corrupted. In contrast, transformer models like Swin-large achieve a better balance between precision and recall, making them more suitable for real-world tilt series processing.

### 4.5 Attention-Based Interpretability

To gain insight into how models differentiate between corrupted and good tilts, we analyzed attention maps generated by Swin_tiny transformer. Figure 5 reveals that attention is distributed differently depending on the type of artifact. The model focuses its attention on key structural regions associated with corruption, such as lamella edge or contamination. Ice reflection regions are well-localized, with model attention concentrated on the high-contrast artifacts. Good tilts and drift artifacts show more diffuse attention patterns, likely contributing to the higher misclassification rates observed in confusion matrices for drift. These results highlight the interpretability advantages of transformer-based models, which not only achieve high F1-scores but also offer meaningful visual cues that provide insight into their decision-making process.

**Figure 5.**
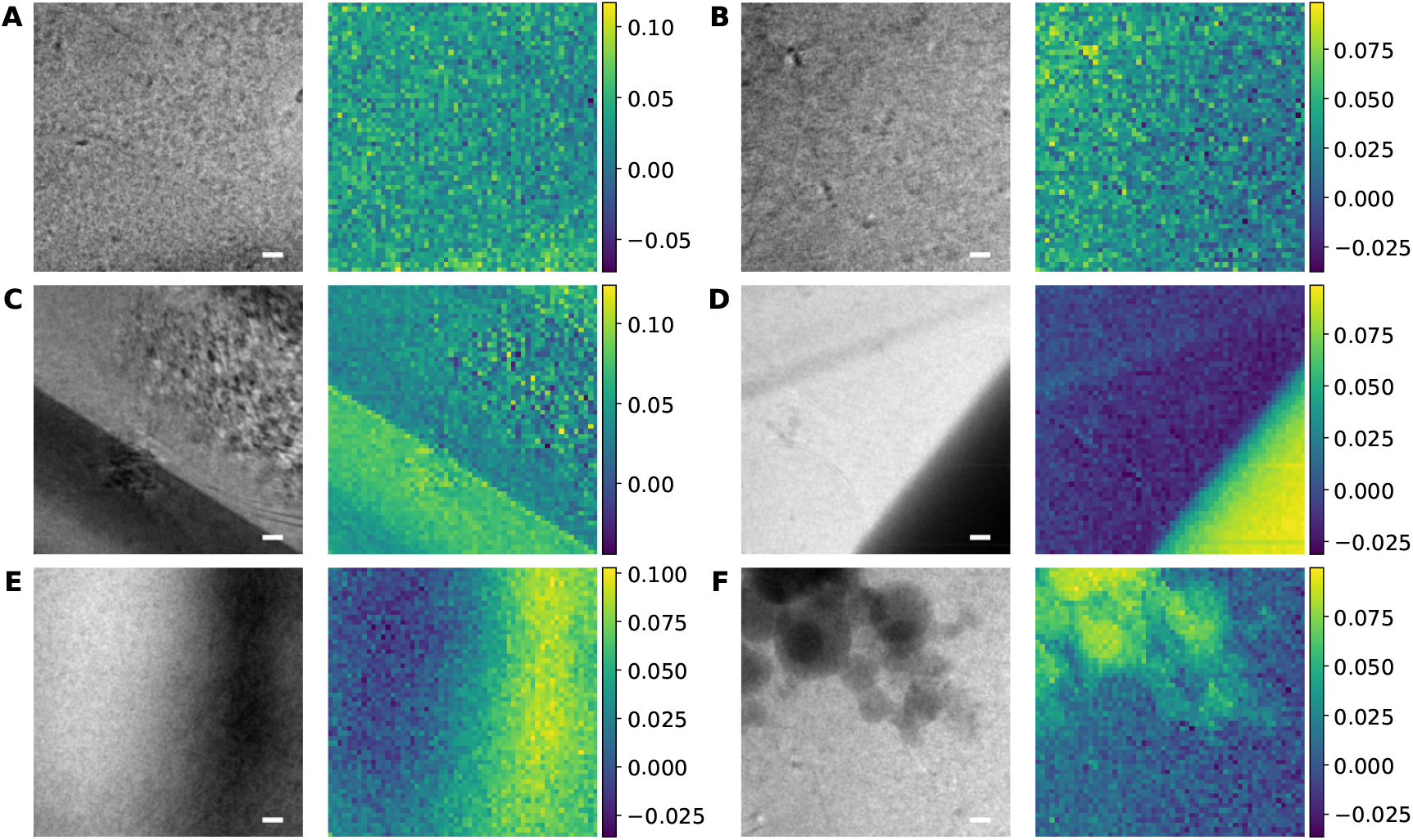
The collection of attention maps for different tilt images. A) Good, (B) Drift, (C) Ice Reflection, (D) Lamella Edge, (E) Thick Lamellae, (F) Contamination. The resolution of the attention map is 56×56 pixels because Swin_tiny transformer is using 4×4 patch embedding on input images of 224×224 pixels. The scale bar is 60 nm.

## 5 Discussion

The results of our study demonstrate that deep learning models can effectively automate tilt cleaning in cryo-ET, reducing the need for manual intervention. Transformer-based models, particularly Swin transformers, consistently performed well in both binary and multiclass classification tasks. ResNet-50 and EfficientNet_b3 remain highly competitive in both settings. EfficientNet_b3 achieved the highest AUC score in binary classification (0.9049), indicating that CNNs remain a viable choice for tilt classification. They are better at recognizing corrupted tilts, but may not have an optimal threshold for making final decisions.

The superior performance of Swin transformers can be attributed to their ability to model long-range dependencies in image data through self-attention mechanisms. As shown in Figure 3, Swin-large and Swin-tiny achieved high AUC-ROC scores, demonstrating robust generalization to unseen data. The self-attention mechanism allows these models to focus on the most relevant regions of an image, as confirmed by the attention maps in Figure 5.

While deep learning models demonstrated strong performance, certain artifact categories remained challenging. In particular, drift and ice reflection distortions were frequently misclassified (Figure 4), especially by transformer models. Drift could potentially be detected by analyzing the Fourier transform of the images, and incorporating such extensions may improve the overall framework performance. Furthermore, integrating generative models could enable the system to learn more nuanced features.

Figure 2B also highlights the impact of dataset size on model performance. Even models fine-tuned with only a small number of additional tilt series showed a marked improvement in F1-score. This suggests that transfer learning from large-scale cryo-ET datasets could further enhance model performance and enable quick adaptation of pre-trained models to a user’s specific cleaning task.

Despite the promising results, our study has some limitations. The models were trained and tested on manually labeled data, which, despite careful curation, may still contain subjective biases. Future work should focus on investigating semi-supervised and self-supervised learning methods to reduce reliance on manually labeled data. We see potential in exploring hybrid architectures that integrate CNN-based local feature extraction with transformer-based global attention mechanisms and deploying real-time inference models that can classify tilts directly during data acquisition.

## 6 Conclusion

In this study, we introduced a deep learning-based method for automated classification of corrupted tilts in cryo-ET. By evaluating 13 deep learning architectures, including CNNs and transformers, we demonstrated that pre-trained models achieve high accuracy in both binary and multiclass tilt cleaning tasks.

Our key findings include:

- Transformer-based architectures, particularly Swin transformers, achieved AUC-ROC scores above 0.96 with the highest F1-scores in our classification test suggesting high correlation with manual annotations. Compared to manual annotation, they correctly identify over 93% of good tilts and more than 70% of corrupted ones.
- CNN models, such as EfficientNet_b3 and ResNet-50, demonstrated higher accuracy in recognizing corrupted tilts. For ResNet-50, this accuracy exceeds 82%, though it may not have an optimal threshold for making final classification decisions.
- Even the addition of a small number of training tilt series markedly improved the performance of pre-trained models, highlighting the potential of transfer learning for cryo-ET applications.
- Our models can process a TS of 50 tilts in under two seconds, and fine-tuning on four TS datasets requires between three and twelve minutes on a single GPU, depending on the model’s size.
- Attention-based visualization revealed that Swin_tiny transformer effectively focuses on relevant structural regions associated with tilt corruption, making it highly interpretable for practical applications.

The proposed automated tilt classification method represents a significant step toward improving the efficiency and reproducibility of cryo-ET data processing. By integrating this approach into existing cryo-ET software suites, researchers can reduce the time spent on manual tilt curation while ensuring consistent and objective tilt selection.

Along with this study, we provide Jupyter Notebook scripts for training and evaluating deep learning models for tilt cleaning. We also include augmentation scripts and scripts for TS-based data splitting into training, validation, and testing subsets, enabling users to train and validate their own models. Additionally, we provide a standalone script for running the cleaning process from the command line. This script generates a cleaned tilt stack along with a PDF highlighting the removed tilts and their predicted probabilities of being corrupted, based on the network’s predictions (see Supplementary Figure 3). In the Data Availability section, we provide our best fine-tuned models used in this study, which can be directly applied and easily integrated into existing workflows.

The introduction of automated tilt removal would have substantial implications for cryo-ET data processing. Current manual tilt selection methods are time-consuming and susceptible to user variability, leading to inconsistencies in data quality. By implementing deep learning-based tilt classification, researchers could standardize the tilt removal process across laboratories and projects, reduce the time spent on manual curation, accelerate the reconstruction pipeline, and improve the reproducibility of tomogram reconstructions by ensuring consistent tilt selection criteria.

## Data Availability

At the time of writing, dataset D2 was published with EMPIAR-12454 and EMPIAR-12457^33^ and D5 with EMPIAR-11845, EMPIAR-11943, and EMPIAR-11944^3^. Fine-tuned binary and multiclass models are publicly available on our ownCloud: https://oc.biophys.mpg.de/owncloud/s/zmMZPr2TEB4Bwda.

## Code Availability

We provide four fine-tuned binary and multiclass models (Swin_tiny, Swin_large, ResNet-50, and EfficientNet_b3), trained on our manually annotated datasets of more than 15,000 tilt images. Users can access these models, along with the provided Python scripts in our GitHub repository https://github.com/turonova/ARCTiC to incorporate them directly into their processing pipeline or to fine-tune them with their own data.

## Acknowledgements

We thank Desislava Glushkova, Patrick C. Hoffmann, Jan Philipp Kreysing, Lung-Yu Liang, Maarten W. Tuijtel, and Huaipeng Xing from the Max Planck Institute of Biophysics for providing data used in the training of our models. We are also grateful to Stefanie Böhm, Jan Philipp Kreysing, Lung-Yu Liang, Maarten Tuijtel, and Martin Beck for their critical reading of the manuscript and valuable discussions. All data presented in this manuscript were acquired at the Central Electron Microscopy Facility of the Max Planck Institute of Biophysics.

## Author information

Conceptualization, Investigation, Methodology, Validation, Formal analysis, Software, Visualization, Writing—original draft, Writing—review & editing: T.M, and B.T., Supervision and funding acquisition: B.T.

