## Supplementary Information for "Automated Removal of Corrupted Tilts in Cryo-Electron Tomography"

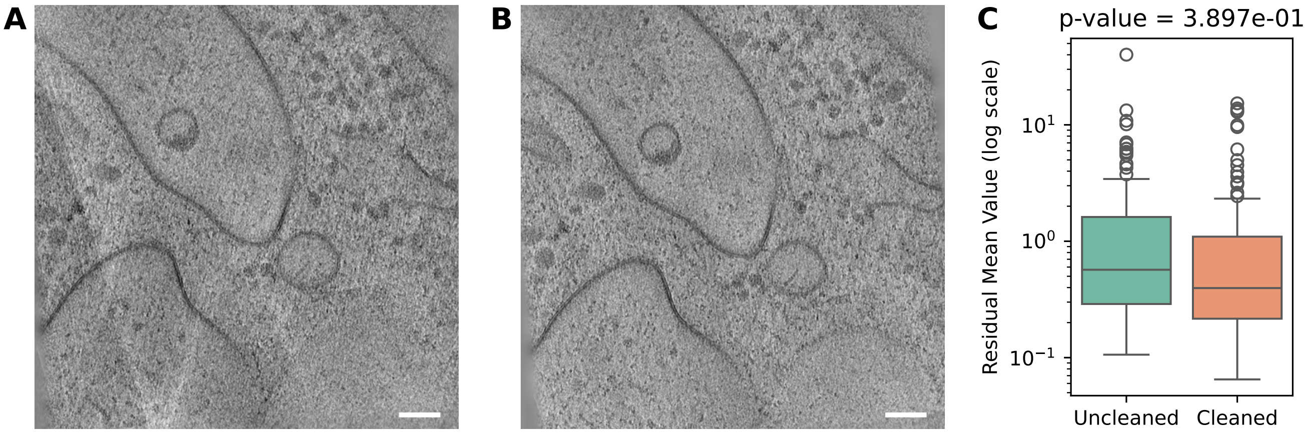


**Supplementary Figure 1.** The effect of cleaning on tomogram reconstruction. (A) The slice of reconstructed tomogram with all tilt series images, including the corrupted ones. (B) The same tomogram slice after the removal of corrupted tilts prior to the tilt series alignment. (C) The summary of all residual mean values of the tilt series alignment from dataset D4 before and after cleaning. The scale bar in (A) and (B) is 40 nm.

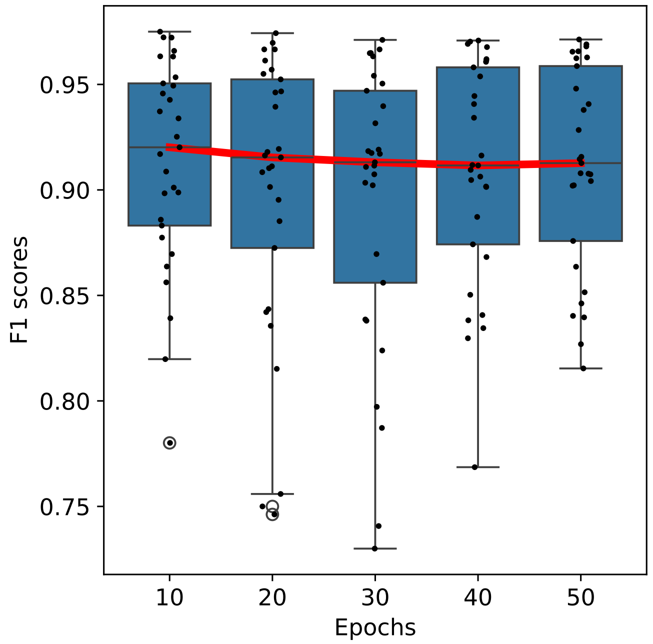


**Supplementary Figure 2.** F1 scores with median trend line across different number of epochs.

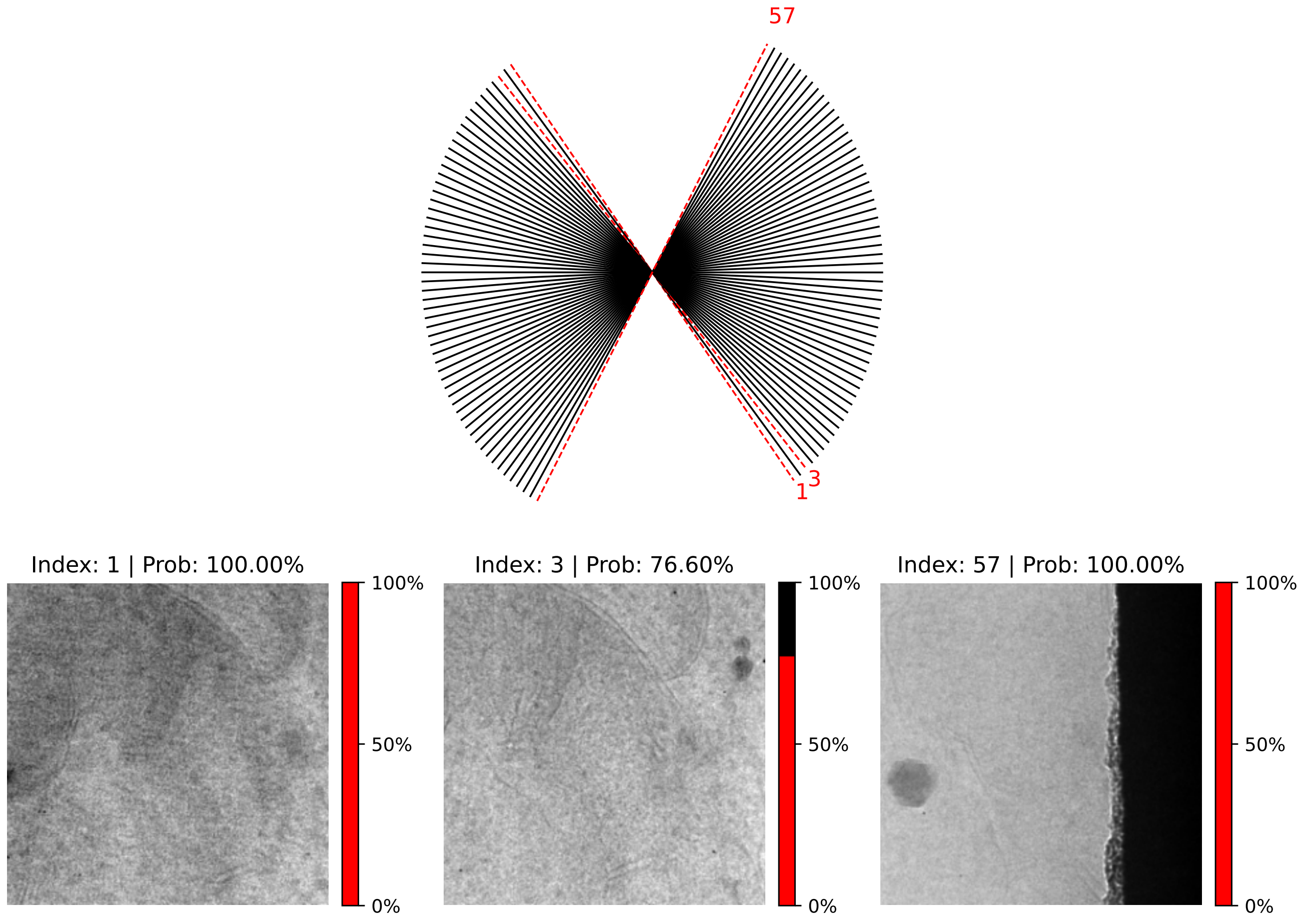


**Supplementary Figure 3.** Visualization of the tilt cleaning process, where the network identifies three corrupted tilt images in the TS and outputs the probabilities for each of them.

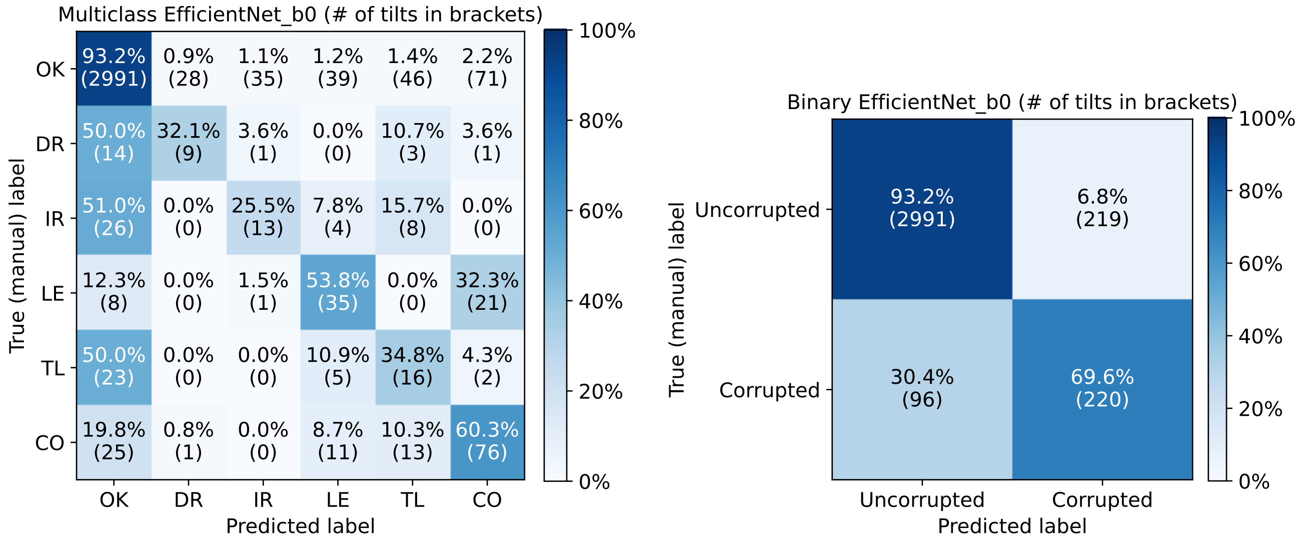


**Supplementary Figure 4.** Confusion matrices for EfficientNet_b0 for multiclass and binary evaluation.

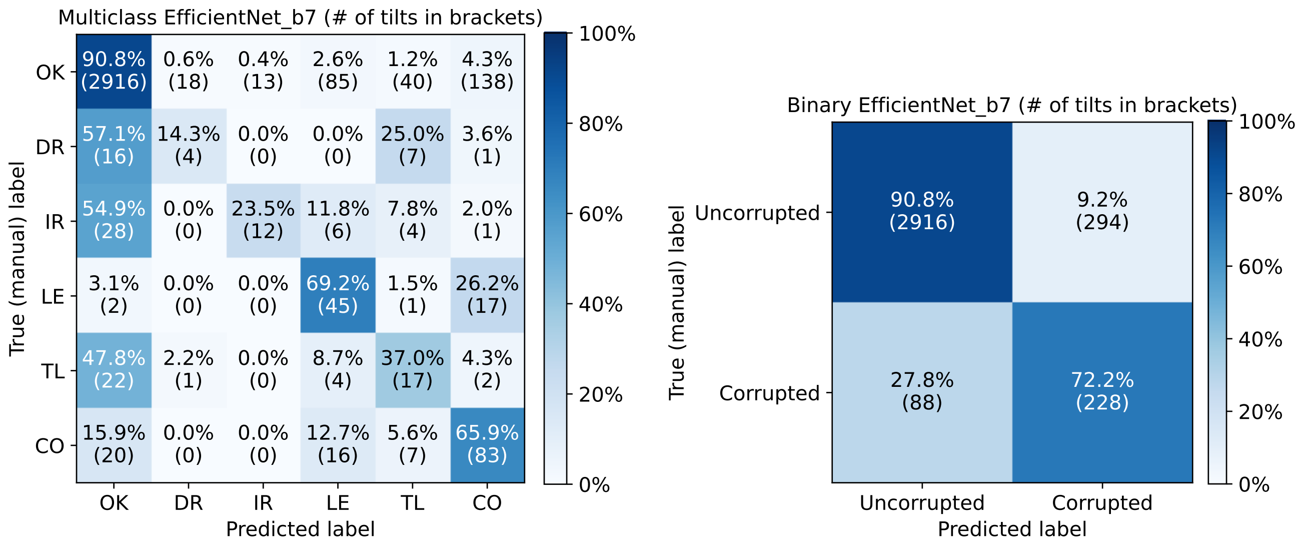


**Supplementary Figure 5.** Confusion matrices for EfficientNet_b7 for multiclass and binary evaluation.

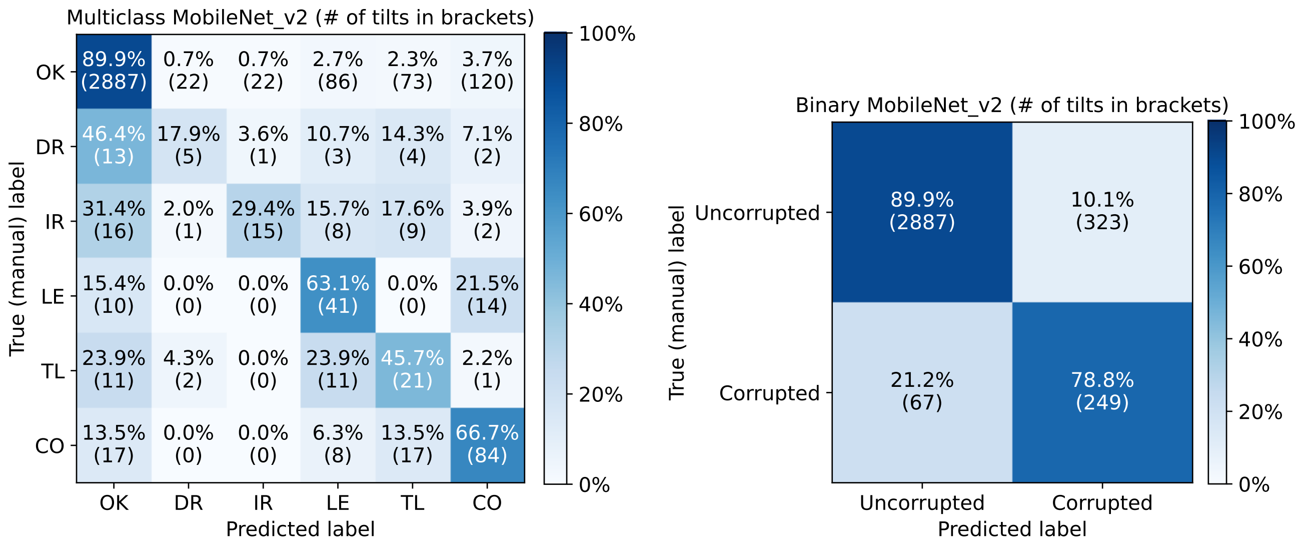


**Supplementary Figure 6.** Confusion matrices for MobileNet-v2 for multiclass and binary evaluation.

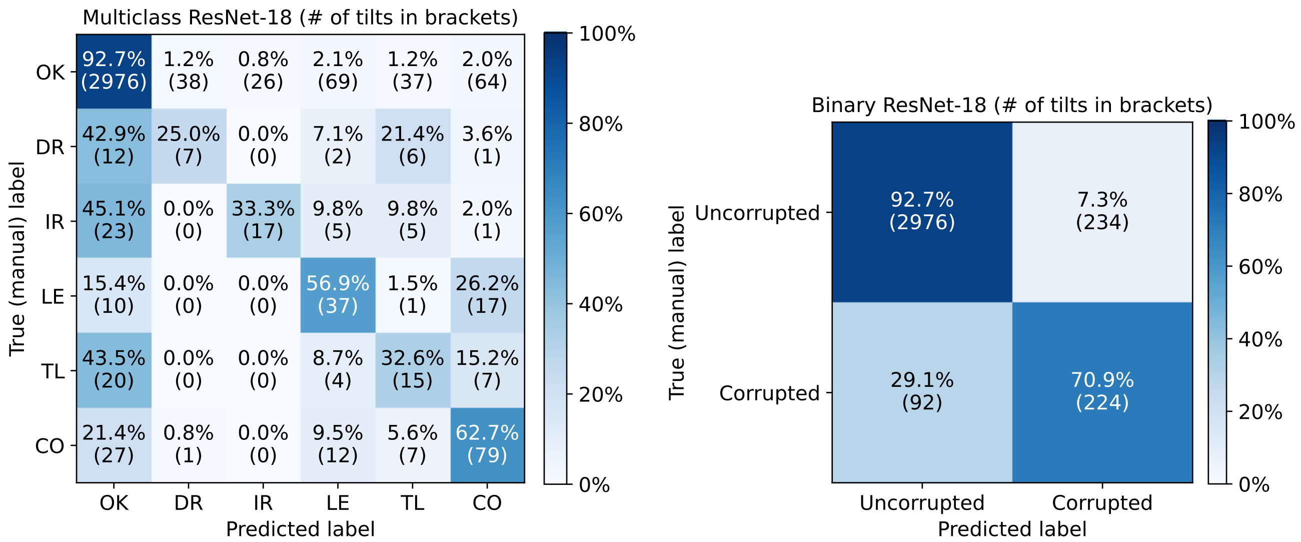


**Supplementary Figure 7.** Confusion matrices for ResNet-18 for multiclass and binary evaluation.

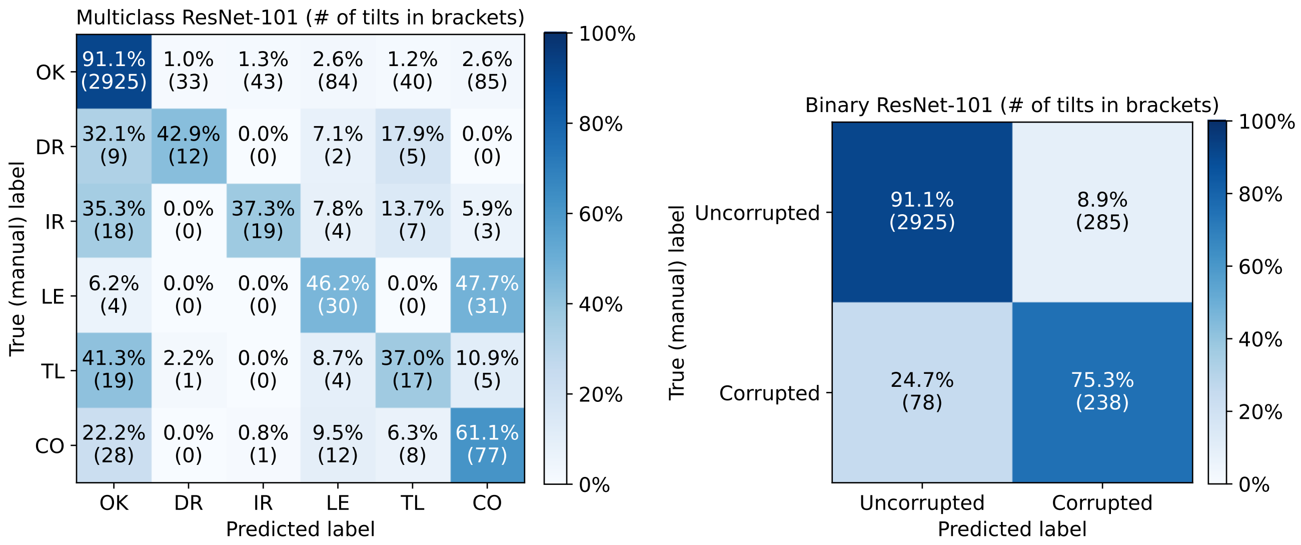


**Supplementary Figure 8.** Confusion matrices for ResNet-101 for multiclass and binary evaluation.

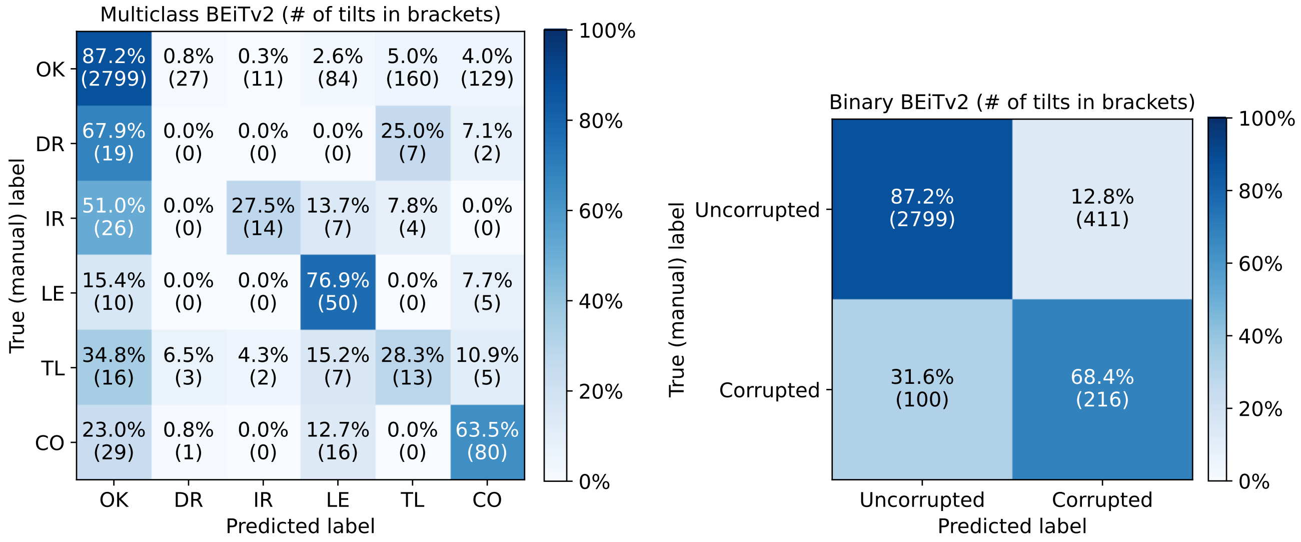


**Supplementary Figure 9.** Confusion matrices for BEiTv2 transformer for multiclass and binary evaluation.

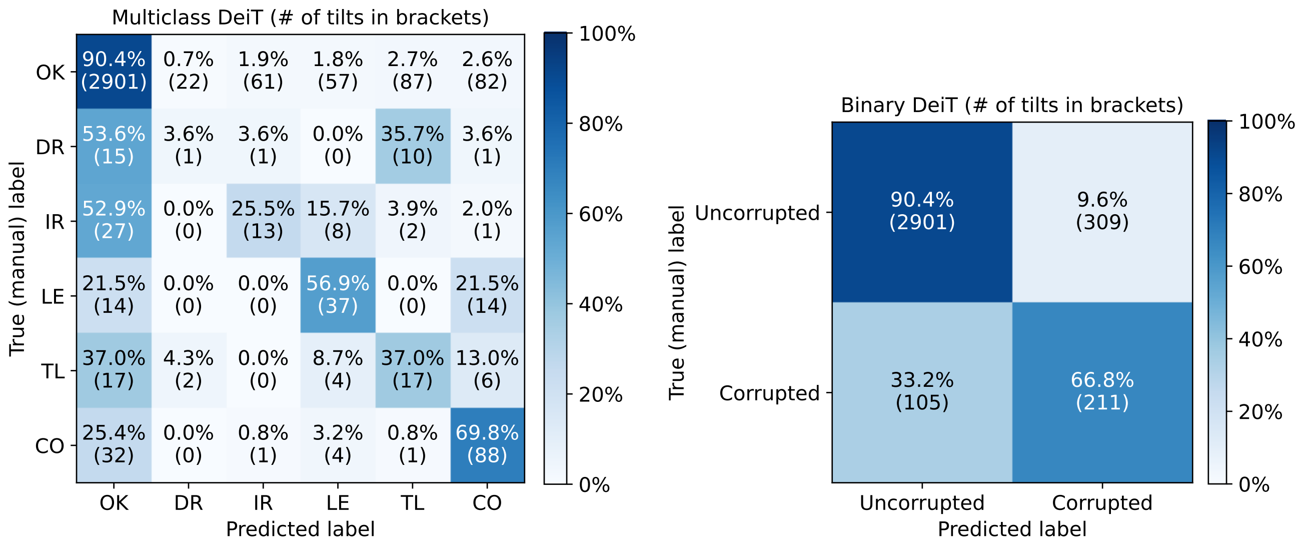


**Supplementary Figure 10.** Confusion matrices for DeiT transformer for multiclass and binary evaluation.

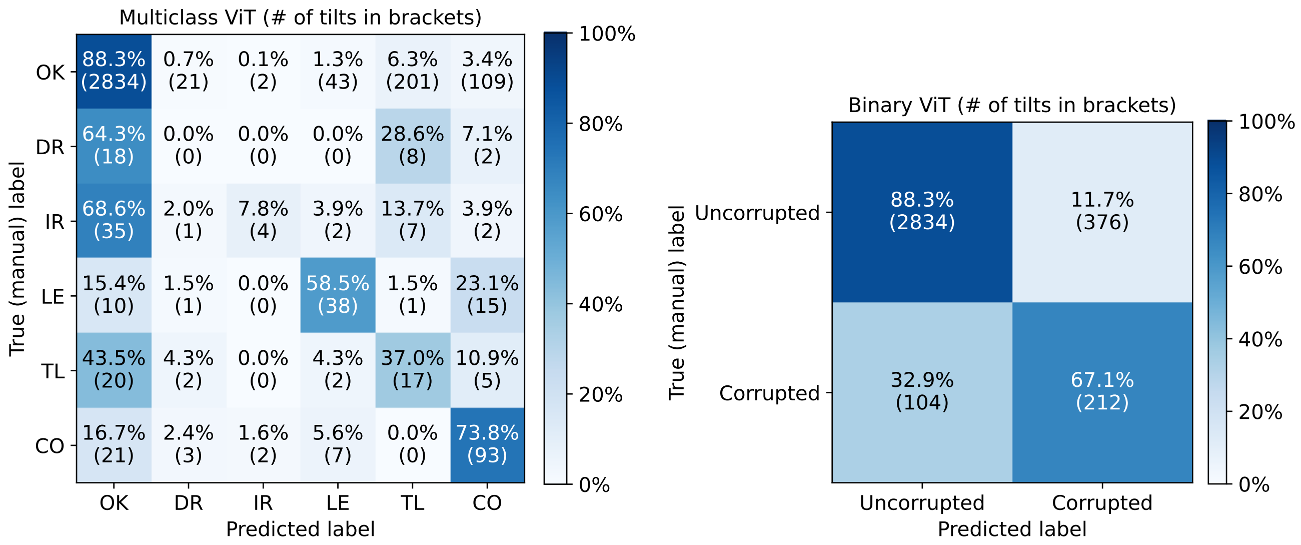


**Supplementary Figure 11.** Confusion matrices for ViT transformer for multiclass and binary evaluation.

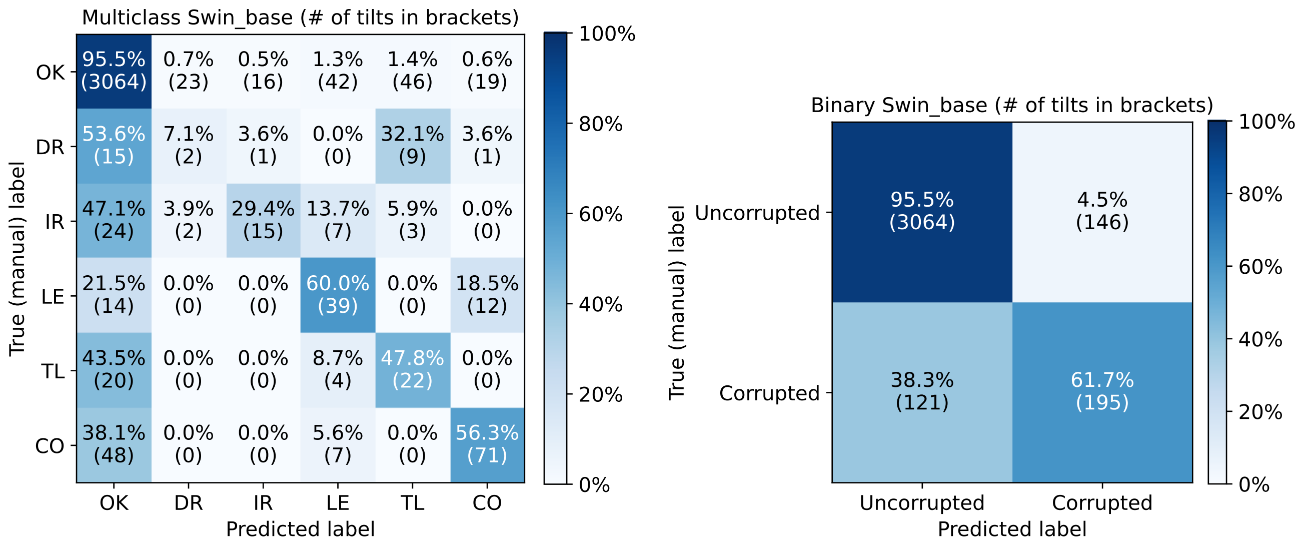


**Supplementary Figure 12.** Confusion matrices for Swin base transformer for multiclass and binary evaluation.

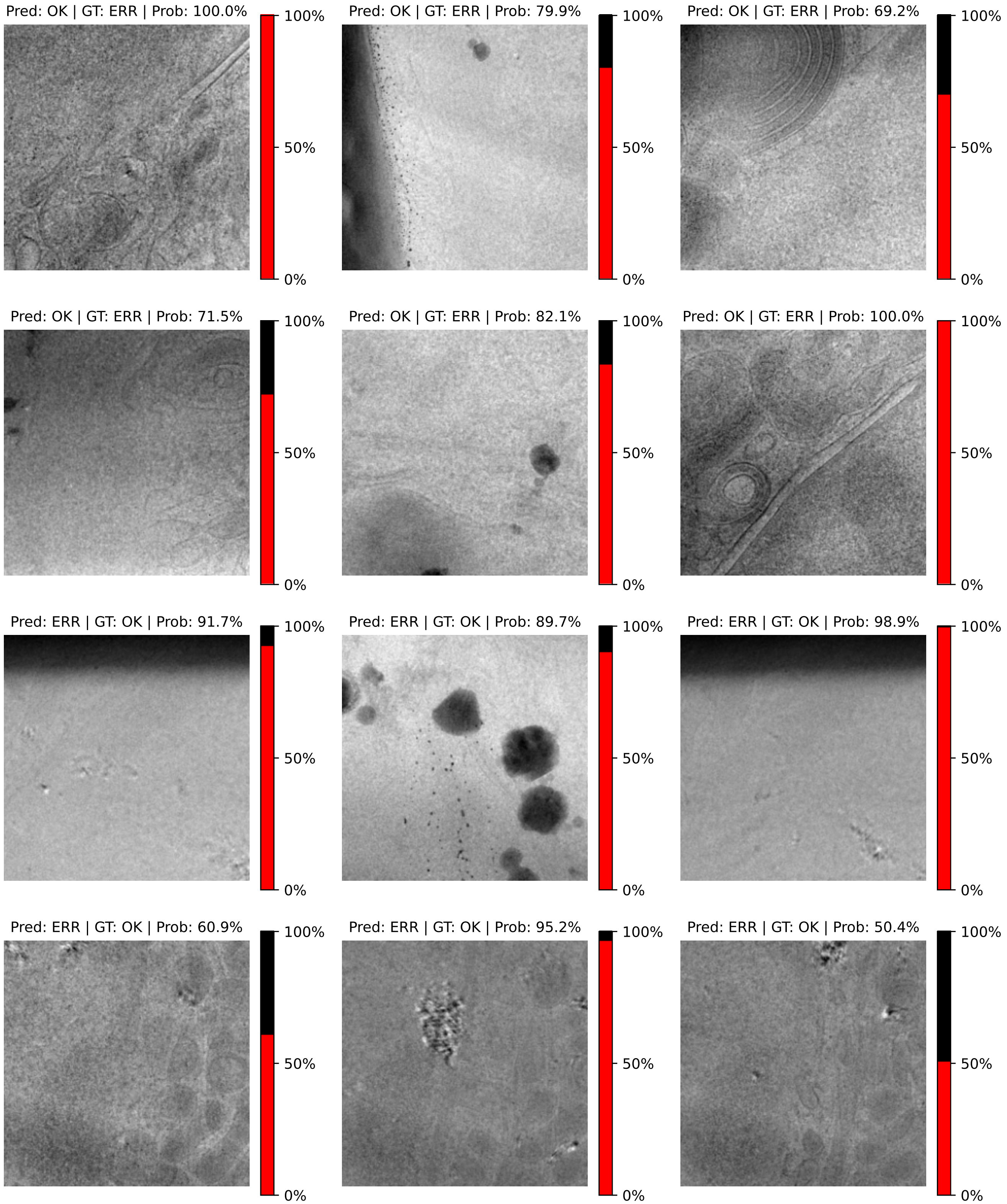


**Supplementary Figure 13.** Examples of misclassified tilt images compared to manually annotated labels provided by users who acquired these images. *Pred* stands for the model's prediction, *GT* represents the ground truth (manual annotations), and *Prob* indicates the certainty of the model's prediction.
